# Age-related changes in brain metabolites underlie slowing of speed-accuracy trade-off

**DOI:** 10.1101/2022.11.02.512356

**Authors:** Lauren Revie, Claudia Metzler-Baddeley

## Abstract

Aging leads to response slowing but the underpinning cognitive and neural mechanisms remain elusive. We studied cognitive components of response speed with a diffusion drift model to estimate nondecision, boundary-separation, and drift-rate contributions to choice reaction times of older (62-80 years) and younger (18-29 years) adults (n=25 each). We characterised age-related differences in the metabolic and microstructural profile of cortical regions and white matter (WM) pathways of visuo-perceptual and attention networks with magnetic resonance spectroscopy and multi-shell diffusion-weighted imaging. Aging was associated with increased non-decision time and boundary-separation, reduced N-acetyl aspartate (NAA) concentrations in anterior cingulate (ACC) and posterior parietal cortices, and reduced WM microstructure in the optic radiation (OR), inferior and superior longitudinal fasciculus (ILF, SLF) and fornix. NAA in ACC and diffusivities in OR and SLF predicted non-decision time, while restricted diffusivity signal fraction in the ILF and fornix, and diffusivity in OR predicted boundary-separation. These results suggest that age-related deterioration of neuronal health and WM microstructure in visuo-perceptual and attention networks contribute to response slowing in aging.

## Introduction

One of the best-established findings in aging research concerns the slowing of response speed and lengthening of the speed accuracy trade off (SAT) as people are getting older (Salthouse, 1979). SAT refers to the trade-off that occurs between responding as timely and as accurately as possible when completing a time-pressured cognitive task. SATs are greater in aging, meaning that older adults adopt a more cautious response strategy favouring correct responses at the cost of slower reaction times (RT) (Starns & Ratcliff, 2010). In contrast, younger adults typically make faster responses which may be at greater risk of error (Der & Deary, 2005). Whilst the exact factors that lead to this strategy change in older adults is yet to be established, it has been attributed to several factors such as a distrust in being able to provide a correct response. One hypothesis proposes that this may be due to age-related deterioration in sensory and perceptual functioning including visual processing. This means that longer perceptual processing time is required to effectively interpret the stimulus, thus increasing overall RTs or increasing the likelihood of an incorrect response if RT is not lengthened (Basak & Verhaeghen, 2011).

Indeed, age-related perceptual decline has been observed at some of the lowest levels of visual processing such as visual contrast, even in those with intact visual acuity and an absence of visual impairment (Delahunt et al., 2008, Elliott et al., 1990, Govenlock et al., 2009). Such low-level impairments in turn can impact mid and higher levels of perceptual and decision-making functions. However, it remains unclear whether cognitive slowing is a cause or consequence of this age-related reduction in visual ability.

The analysis of mean RTs or SAT alone cannot inform us about the cognitive elements that contribute to age-related slowing, but they can be modelled to offer a solution. The drift diffusion model (DDM) can be applied to RT data from choice RT tasks such as the flanker task (Eriksen & Eriksen, 1974) to assess different hypothetical processing stages involved in decision making, including the contribution of perceptual/and or motor processes (Ratcliff, 1978; Ratcliff & Rouder, 1998). The Eriksen flanker task is a classic response inhibition task that involves the presentation of a target arrow flanked by distractor arrows, which are either congruent with the directional response to the target, i.e., a left or right key press, incongruent (pointing into the opposite direction), or neutral. Estimating the contribution of different decision-making elements to older compared to younger adults’ performance in such a flanker task allows an insight into the sub-processes impacted in aging. This may help tease apart the contributions of bottom-up perceptual and/or top-down decision-making elements, such as the evaluation of the quality of processing, to age-related response slowing. The DDM is a sequential-sampling model (Ratcliff, 1978; Ratcliff & Rouder, 1998) which in simplest terms allows the estimation of the processing time for each element within the RT. The model assumes that information which drives a decision is accumulated over time until it reaches one of two response boundaries to form the final response. According to the DDM, overall processing of a decision is segmented into components which contribute to the ultimate overall performance: first, components of processing which do not include active decision, such as perceptual encoding and response execution, represented by ‘non-decision time (t)’, second, a criteria threshold that information accumulates to, represented by ‘boundary separation (a)’, and third, the rate at which information accumulates towards a decision, represented by ‘drift rate (v)’. The DDM assumes that the accumulation of information (drift rate) during a decision process is not constant but varies over time, represented by (s) (Figure 1).

**Figure 1.**
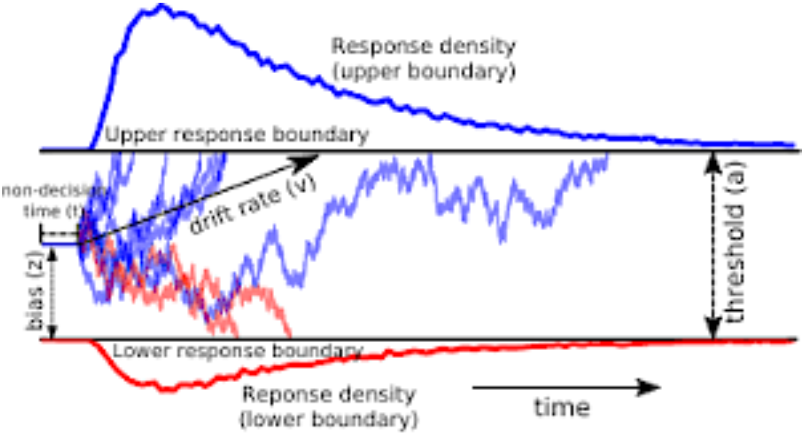
The drift diffusion model (DDM) of response time. Red and blue lines denote different responses in a 2-choice task (for example, left or right). Reaction times (RT) are fit to the DDM to return an estimate of nondecision time (t) reflecting perceptual and motor processing time, boundary separation value or threshold (a), representing the level at which information must reach to trigger a response and drift rate (v), representing the efficiency of the drift process. Adapted from Zhang & Rowe (2014).

Older adults consistently show increased ‘boundary separation values’, which refers to the threshold stimulus processing must reach before a response decision can be made (Rabbitt, 1979). This indicates that older adults require increased stimulus processing time in order to make a response. In addition, a lengthening of non-decision time, i.e., the perceptual and motor element of the decision, has also been observed in older adults (Rabbitt, 1979). However, it remains unknown whether this lengthening of nondecision time is due to impaired bottom-up low-level visual functioning, slowed motor responses or both, and whether visual perceptual difficulties contribute to age-related increases in boundary separation.

Furthermore, little is known about the neural substrates that underpin the age-related changes in different decision-making components. Only a few studies have explored correlations between age-related differences in DDM parameters and in brain structure and function in the absence of retinal pathology (Forstmann et al., 2011, Monge et al., 2017, Yang et al., 2015). These studies have identified correlations between age-related differences in DDM parameters and white matter microstructure such as negative correlations between boundary separation and fractional anisotropy in fronto-cortical tracts (Forstmann et al., 2011), as well as positive correlations between non-decision time and radial diffusivity in the corpus callosum (Yang et al., 2015). Furthermore, differences in functional MRI activity of fronto-parietal regions were linked to age-related variability in drift rate (Madden et al., 2010). These findings demonstrate associations between age-related structural and functional changes of connections within fronto-parietal decision-making networks and age-related slowing of DDM components. This accords with evidence of correlations between fronto-parietal white matter impairments and age-related slowing in simple and choice RT tasks and in SAT (Bugg et al., 2006, Jackson et al., 2012, Madden et al., 2004, Monge et al., 2017).

These white matter impairments in older age are thought to occur primarily due to accumulating damage and loss of myelin, the fatty substance that insulates axons and facilitates saltatory conduction and is hence crucial for neuronal health and efficient information transfer across the brain (Bartzokis et al., 2010, Fields, 2005). However, neurochemical metabolites such as N-acetyl-aspartate (NAA) and myoinositol are pivotal in the production and maintenance of myelin and neuronal health (Glanville et al., 1989, Rae, 2014). Moreover, accumulating evidence from magnetic resonance spectroscopy (MRS) studies suggest that their concentration changes in the aging brain and that such changes may underpin age-related slowing (Simmonite et al., 2019). Despite this evidence, the neurochemical basis for age-related changes in SAT are yet to be investigated. MRS is a non-invasive MRI technique which is employed to quantify metabolites *in vivo* by acquiring frequency spectra to assess the biochemical composition of tissues in a selected brain region. Whilst MRS can be conducted as a whole-brain technique, spectra are more commonly assessed in specific regions of interest (ROIs) or voxels, to acquire a level of signal to noise which can be accurately quantified for multiple metabolites. Most commonly quantified metabolites are N-acetyl aspartate (NAA), creatine (Cr), choline (Cho), myoinositol (mI), glutamate/glutamine (Glx) and γ-aminobutyric acid (GABA). NAA and creatine are known to play a role in energy metabolism in neuronal mitochondria (Lu et al., 2004) and glia respectively (Urenjak et al., 1993), where choline is involved in processes such as inflammation and demyelination (Zeisel & da Costa, 2009). In addition, myoinositol has been proposed as a marker of glial cell proliferation (Glanville et al., 1989), and Glx and GABA are the brain’s major excitatory and inhibitory neurotransmitters respectively (Stagg & Rothman, 2013). These metabolites are all shown to vary with aging and are linked cellular and microstructure activity associated with brain aging.

Healthy aging has generally been associated with increases in choline, myoinositol, and creatine but with reductions in NAA, Glutamine (GLx) and GABA (Haga et al., 2009). Such patterns have been observed globally in the brain including in the occipital, frontal, and anterior and posterior cingulate cortices (Chiu et al., 2014; Gruber et al., 2008; Haga et al., 2009; Gao et al., 2013; Marjańska et al., 2017; Pitchaimuthu et al., 2017; Porges et al., 2017; Simmonite et al., 2019) although some studies did not find any age-related changes (see eg. Ferguson et al., 2002; Harada et al., 2001; Marjańska et al., 2017; Ross et al., 2005).

Whilst no studies have specifically investigated the relationships between these age-related changes in brain metabolites and SATs, one study found a link between an age-related decline in fluid processing ability (including speed of processing) and reductions in occipital GABA (Simmonite et al., 2019). This suggests that a reduction in inhibitory neurochemicals in brain areas responsible for visual functioning may contribute to the age-related changes observed in response slowing of other fluid functions.

The integrity and functioning of neural networks at a cellular level are key to their efficient processing, including speed of processing and visual abilities. In particular, the structure and integrity of white matter pathways is integral to the communication between and within these networks, and declines in white matter microstructure have been linked to reduced processing speed in aging (Forstmann et al., 2011). Further, it stands to reason that the neurochemical processes which regulate white matter myelin maintenance and repair may also provide important information regarding the basis for changes in processing speed. Moreover, these neurochemicals moderate energy usage as well as inhibitory and excitatory control within neural networks, and thus play a key role in information processing.

However, while age-related differences in white matter microstructure were found to be related to differences in DDM parameters, and neurochemicals are thought to play a pivotal role in the maintenance of myelin microstructure, it remains unknown whether metabolic changes in visuo-perception and attention networks contribute to the slowing of decision-making processes with age.

Thus, the present study used MRS and multi-shell diffusion-weighted MRI to investigate the metabolic and microstructural substrates of age-related differences in DDM parameters that may contribute to cognitive slowing in aging. Concentrations of metabolites reflective of neuronal healthy (NAA), inhibitory or excitatory functioning (Glx and GABA), energy production and usage (Glx, creatine, choline), and glial functioning (mL) were measured in three cortical ROIs, i.e., the anterior cingulate cortex (ACC), the posterior parietal cortex (PPC), and the occipital cortex (OCC). These ROIs were selected as they form key regions of visual perception and attention networks that mediate bottom-up and top-down processing streams. Further, the microstructure of key white matter pathways that connect these visual and attention network regions and are known to be involved in top-down and/or bottom-up processing and attentional functioning, i.e., the superior longitudinal fasciculus (SLF), the inferior longitudinal fasciculus (ILF), the optic radiation, and the fornix was studied. White matter microstructure of these tracts was characterised with metrics from the Composite Hindered and Restricted Model of Diffusion (CHARMED) (Assaf & Basser, 2004) that allows the separation of intra- and extra-axonal contribution to the diffusion signal. CHARMED was employed in this investigation as it provides the restricted diffusion signal fraction (FR), a proxy index for axonal density, as well as diffusion tensor-based metrics of fractional anisotropy (FA), mean diffusivity (MD, axial diffusivity (L1) and radial diffusivity (RD). FR is a valuable metric to quantify in this context, as it has been shown to be more sensitive than DTI indices and has been suggested as a potential biomarker for axonal microstructure changes (De Santis et al., 2017) which are well-established in aging.

These metabolic and microstructural measurements from anterior, parietal and occipital cortices and their white matter connections were acquired from healthy young (n = 25, age range = 18-29 years) and older participants (N = 25, age range = 62-80 years) to study the contributions of bottom-up visual perceptual and top-down decision-making processes to age-related differences in DDM parameters and SAT. Participants performed a modified version of the Attention Network Test (ANT) that involves a choice flanker task (Fan et al., 2002) and SAT and DDM parameters were calculated from participants’RTs in this task.

In this way we were able to characterise age-related metabolic differences in key structures of the visual and attentional networks and to study how these differences were related to differences in SAT and DDM parameters. We hypothesised that aging would be associated with reductions in NAA, GABA, and Glx and increases in choline, myoinositol and glutamate in OCC, ACC and PPC as well as with reductions of FA and FR and increases in MD, RD and L1 in SLF, ILF, optic radiation and fornix. We further hypothesised that age-related metabolic and microstructural differences in lower-level perceptual and sensory processing areas (occipital lobe, optic radiation, ILF) would be associated with differences in nondecision time, while those in SLF, ACC and PPC would be associated with differences in boundary separation and drift time due to their roles in top-down processes.

## Methods

### Participants

Participants were recruited from the School of Psychology community participant panel at Cardiff University and consisted of younger (aged 18-29) and older (aged 62-80) adults. Twenty-six participants were recruited into each group, all of whom provided informed written consent prior to taking part in the study in accordance with the Declaration of Helsinki (Cardiff University School of Psychology Ethics committee reference 18.06.12.5313; NHS Research Ethics committee REC reference 18/WA/ 0153). All participants were cognitively healthy, i.e., had a Montreal Cognitive Assessment (MOCA) score >= 26. Participants also completed MRI screening prior to the study, excluding any participants with MRI contraindications such as metallic or electronic bodily implants, some dental work and some tattoos, subject to radiographer assessment. Individuals with visual impairments, such as visual field loss or glaucoma were also excluded from the study. Table 1 summarises participants’ demographic information as well as their mean performance on cognitive and visual screening tasks.

**Table 1.** Demographic and baseline cognitive scores for younger and older adults. Mean and standard deviation (SD) for younger and older adults’ performance. MOCA = Montreal cognitive assessment, TOPF = test of premorbid functioning.

|  | Younger Mean (SD) | Older Mean (SD) |
| --- | --- | --- |
| Age | 21.65 (2.76) | 68.46 (5.91) |
| Gender | Male (12) Female (14) | Male (9) Female (17) |
| Handedness | Left (1) Right (28) | Left (3) Right (22) |
| Years of education | 14.95 (1.89) | 15.12 (2.40) |
| MOCA score | 28.89 (1.25) | 29.04 (1.14) |
| Visual acuity (Snellen fraction) | 1.91 (0.18) | 1.66 (0.48) |
| TOPF-UK score | 60 (6.43) | 62.52 (13.3) |

### Materials & Procedure

#### Cognitive and visual testing

Testing was conducted at Cardiff University Brain Research Imaging Centre (CUBRIC), during one visit lasting approximately 2 hours. Participants completed a visual acuity task and flanker task on a computer which is described in detail below. The task was presented on a 15” screen (1440 × 900 native resolution) and responses were recorded using a wireless keypad. The flanker task was written by LR using PsychoPy psychophysics software (Peirce, 2009) for Python (v1.85.6) following the original methodology of the Attention Network Task (ANT) (Fan et al., 2002) unless otherwise stated. Visual acuity was assessed using the Freiburg Visual Acuity and Contrast Test (FRACT; Bach, 1996). Participants viewed the screen from a distance of 2m (as recommended by test manufacturers) and responded to circular stimuli, where the target was a ‘gap’ in the circle. Stimuli was reduced in size for each correct trial to achieve a Snellen fraction measure of visual acuity.

Reaction times (RTs) were recorded using a modified ANT flanker task (Fan et al., 2002), and speed accuracy trade-off and DDM parameters were calculated using these RTs. The modified ANT stimuli consisted of five horizontal arrows presented on the screen in which participants were instructed to attend to the central arrow as the target. Central arrows were flanked by horizontal lines (neutral condition), arrows facing in different directions to the target (incongruent condition), or arrows facing in the same direction as the target (congruent condition). During this version of the ANT, stimuli were presented in the same central position on the screen following the presentation of a fixation cross. Participants viewed the screen from a seated position, 400mm from the computer screen. Participants were instructed to maintain focus on the central fixation point of the screen and respond as quickly and accurately as possible. In accordance with the original study (Fan et al., 2002), these stimuli subtended 3.08°of visual angle. Fixations were presented for a random variable length of time between 400-1600ms and target stimuli were presented for a maximum of 1700ms. Participants completed 96 trials (32 trials per condition) in each block, for a duration of 5 blocks. Between blocks, participants were instructed to rest for 30 seconds, before being given a 5 second count-down into the following block. The entire task totalled 480 trials and took approximately 12-15 minutes to complete.

#### Drift diffusion modelling (DDM)

DDM parameters were calculated using the EZ DDM model (Wagemakers et al., 2007) which was incorporated into an in-house R based custom script. Raw RT and accuracy data for each participant for congruent, neutral and incongruent trial conditions were input into the script in R Studio (v 1.1.463). The script first calculated means and variances of correct RTs. Incorrect trials were not included in the remainder of the analysis (average retained trials = 469). Following this, the script calculated DDM parameters using the equation provided in Wagenmakers et al., (2007) under the assumption that trial-to-trial variability was zero and the starting point of each decision process was equidistant from the response boundaries (Schmiedek et al., 2007). This resulted in average estimates for non-decision, boundary separation and drift rate for each participant. Details of the mathematical basis for the EZ model can be found in Wagenmakers, Van der Maas & Grasman (2007).

#### Magnetic Resonance Imaging (MR) Imaging and Spectroscopy

##### MR data acquisition

All MR data were acquired on a Siemens 3 Tesla (T) Magnetom Prisma MR system (Siemens Healthcare GmbH, Erlangen) fitted with a 32-channel receiver head coil at CUBRIC. A 3D, T1-weighted magnetization prepared rapid gradient-echo (MP-RAGE) structural scan was acquired for each participant (TE/TR = 3.06/2250ms, TI = 850ms, flip angle = 9deg, FOV = 256mm, 1 × 1 × 1mm resolution, acquisition time = ~6min). The MPRAGE was used as anatomical reference for the placement of MRS region of interest voxels.

MRS was used to acquire frequency spectra to quantify metabolites of Glx, GABA, NAA, choline, creatine and myoinositol. Single voxel proton spectra were obtained from voxels of interest placed in the occipital cortex (OCC, voxel measuring 30 × 30 × 30 mm^3^), the posterior parietal cortex (PPC, voxel measuring 30 × 30 × 30 mm^3^) and the anterior cingulate cortex (ACC, voxel measuring 27 × 30 × 45mm). The OCC voxel was placed above the tentorium cerebelli, avoiding scalp tissue in order to prevent lipid contamination to the spectra. The PPC voxel was placed with the posterior edge against the parietooccipital sulcus, and the ventral edge of the voxel above and parallel to the splenium. Finally, the ACC was placed directly dorsal and parallel to the genu of the corpus callosum. In each voxel, a spectral editing acquisition (MEGA-PRESS, Mescher et al., 1998) was performed, involving applying an additional pulse symmetrically about water resonance, providing ‘on’ and ‘off’ editing pulses which allow for the subtraction of peaks which may mask GABA in the spectra (TE/TR = 68/2000ms, 168 averages, acquisition time = ~12 min per voxel). Manual shimming was performed before all MRS scans to ensure water-line width of 20Hz or lower, in order to obtain accurate peaks in the spectra (Figure 2).

**Figure 2.**
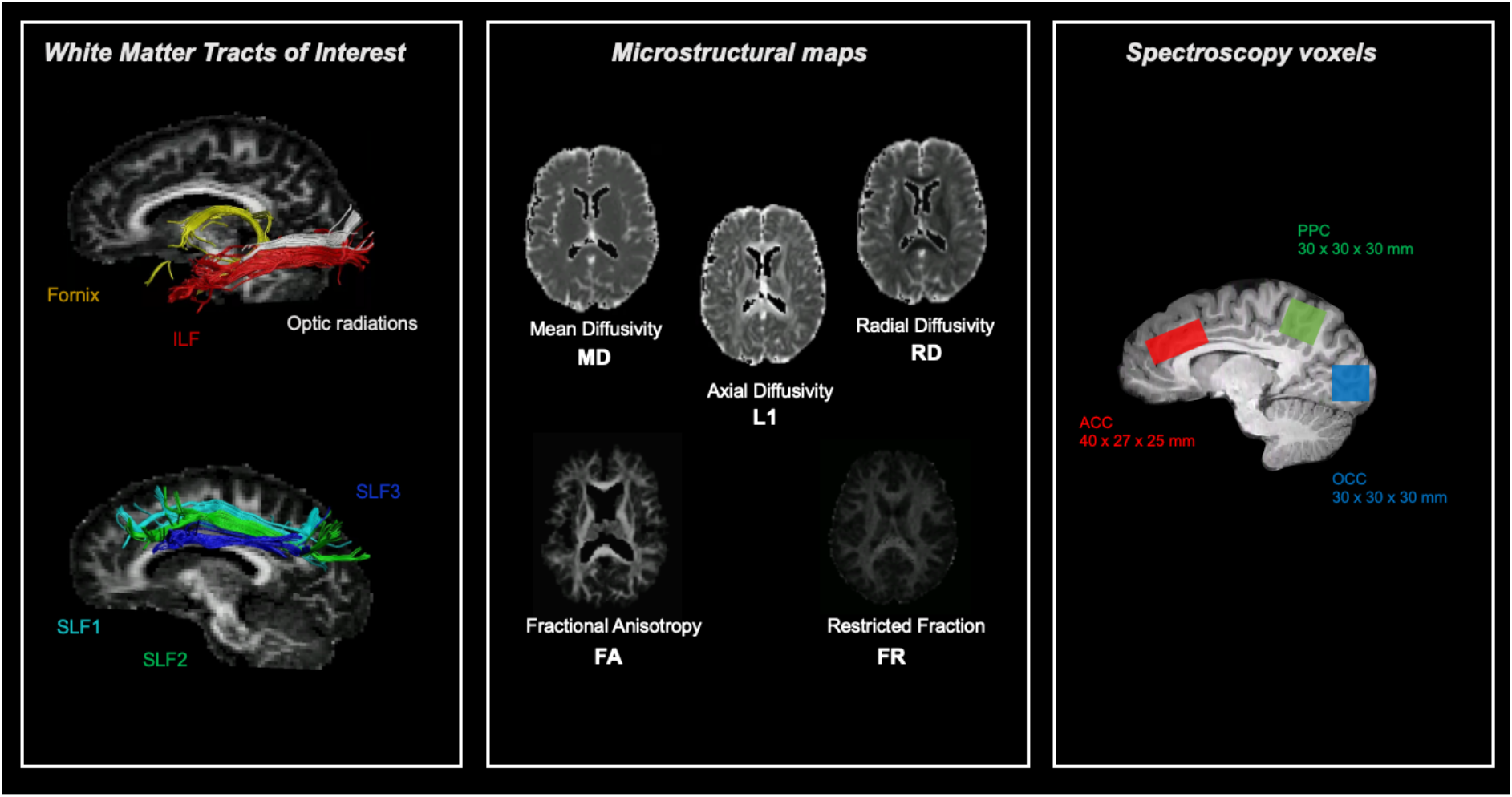
White matter tracts, microstructural maps and location of spectroscopy voxels for measurements of interest. ACC = anterior cingulate cortex, ILF= inferior longitudinal fasciculus, PPC = posterior parietal cortex, OCC= occipital cortex, SLF = superior longitudinal fasciculus.

A multi-shell diffusion MRI sequence was also conducted using a high angular resolution diffusion (HARDI) weighted echo-planar imaging (EPI) sequence (TE/TR = 73/ 4100ms, FOV = 220×220mm, isotropic voxel size 2mm^3^, 66 slices, slice thickness 2mm, acquisition time ~15 min, 2 × 2 × 2mm resolution). Five diffusion weightings were applied along gradient directions: b = 200 s/mm^2^ (20 directions), b= 500 s/mm^2^ (20 directions) b = 1200 s/mm^2^ (30 directions), b=2400 s/mm^2^ (61 directions), b =4000 s/mm^2^ (61 directions). 12 unweighted (b0) volumes were acquired, interspersed throughout diffusion-weighted scans. In addition, a diffusion reference sequence was acquired for later blip-up blip-down analysis to correct for EPI distortion (Bodammer et al., 2004) in which a diffusion weighting of b=1200 s/mm^2^, and 12 un-weighted (b0) images were acquired interspersed throughout the sequence (Figure 1). Multi-shell diffusion weighted imaging data were acquired to fit the diffusion tensor and the Composite Hindered and Restricted Model of diffusion (CHARMED) (Assaf & Basser, 2005) to gain microstructural maps of fractional anisotropy (FA), mean diffusivity (MD), radial diffusivity (RD), axial diffusivity (L1), and restricted signal fraction (FR).

##### MR analysis

MRS data were analysed using Totally Automatic Robust Quantification in NMR (TARQUIN) version 4.3.11 (Reynolds, Wilson, Peet & Arvanitis, 2006) in order to determine estimated concentrations of other metabolites of interest (Choline, NAA, Glx, Creatine, Myoinositol). To ensure data quality, metabolites were excluded if the Cramer Rao Lower Bound (CRLB) was above 20% as recommended (Stagg & Rothman, 2013). MEGA-PRESS data were analysed using GANNET (GABA-MRS Analysis Tool) version 3.0 (Edden et al., 2014). Estimated metabolite values were corrected to account for cerebrospinal fluid (CSF) voxel fraction, and water reference signal was corrected to account for differing water content of CSF, grey matter and white matter. All metabolites were quantified using water as a concentration reference and were expressed as concentration in millimoles per unit (mM).

Two-shell HARDI data were split by b-value (b=1200, and b=2400 s/mm^2^) and were corrected for distortions and artifacts using a custom in-house pipeline in MATLAB and Explore DTI (Leemans et al., 2009). Correction for echo planar imaging distortions was carried out by using interleaved blip-up, blip-down images. Tensor fitting was conducted on the b=1200 s/mm^2^ data, and the two compartment ‘free water elimination’ (FWE) procedure was applied to improve reconstruction of white matter tracks close to ventricles (Pasternak et al., 2009) and to account for partial volume contamination due to CSF which is particularly apparent in older age (Metzler-Baddeley et al., 2012). Data were fit to the CHARMED model (Assaf & Basser, 2005) which involved the correction of motion and distortion artefacts with the extrapolation method of Ben-Amitay et al., (2012). The number of distinct fibre populations in each voxel (1, 2 or 3) was determined using a model selection approach (De Santis et al., 2014) and FR maps (Assaf & Basser, 2005) were then extracted by fitting the CHARMED model to the DWI data, with an in-house script. This resulted in FA, MD, RD, L1 and FR maps.

Whole brain tractography was then performed with the dampened Richardson-Lucy (dRL) spherical deconvolution method (Dell’Acqua et al., 2010). Tractography was performed on the b=2400 s/mm^2^ data to provide better estimation of fibre orientation (Vettel et al., 2012). The dRL algorithm extracted peaks in the fibre orientation density function (fODF) in each voxel using a step size of 0.5mm. Streamlines were terminated if directionality of the path changed by more than 45 degrees using standardised inhouse processing pipeline at CUBRIC. Manual fibre reconstructions were performed in ExploreDTI v4.8.3 (Leemans et al., 2009). Tracts of interest were manually drawn on direction encoded colour FA maps in native space. ILF reconstruction was obtained according to protocol by Hodgetts et al. (2015) and Wakana et al. (2007). The SLF was subdivided into it’s three components, the SLF1, 2 and 3 which were delineated according to protocol by De Schotten et al., (2011). The SLF was subdivided based on the distinct contributions of each tract to different functions of sensory processing; the SLF 1 being associated with spatial functions and goal-directed attention (De Schotten et al., 2011; Parlatini et al., 2017), the SLF2 being associated with orienting attention and integration of dorsal and ventral attention networks (Nakajima et al., 2019), and the SLF 3 being associated with reorienting of spatial attention (De Schotten et al., 2011; Nakajima et al., 2019). The fornix was reconstructed by locating the body of the fornix bundle according to Metzler-Baddeley et al. (2011), and the optic radiation was delineated by placing a seed region on the white matter of the optic radiation lateral to the lateral geniculate nucleus in the axial plane (Thompson et al., 2014) (Figure 2).

### Statistical Analysis

Speed accuracy trade-off (SAT) was calculated from RTs using the linear integrated speed accuracy score (LISAS; Vandierendonck, 2017) method, which combines RT and proportion of error in a linear manner, according to the formula (Equation 1).

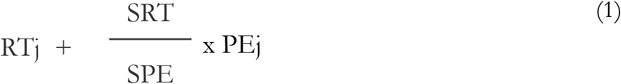

Where RTj is the mean RT, PEj is the proportion of errors, SRT is the participants’ overall RT standard deviation, and SPE is the participants’ overall standard deviation for the proportion of errors.

Tractography outcome measures (FA, MD, RD, L1, FR) and metabolite outcome measures (GABA, NAA, Glx, Myoinositol, Choline, Creatine) were compared between older and younger control groups by conducting non-parametric Mann Whitney U tests. Non-parametric tests were employed as Kolmogorov-Smirnov testing revealed non-normal data distributions of metabolites and microstructural values. Group differences in DDM parameters, SAT, accuracy, RT and variance were assessed using MANOVA with post-hoc tests. Multiple comparisons were corrected for False Discovery Rate (FDR) to mitigate the likelihood of Type 1 error by employing the Benjamini-Hochberg procedure at 5% (Benjamini & Hochberg, 1995).

Two separate linear regression models were carried out for each EZ DDM parameter as dependent variable, one to test for the effects of grey matter metabolites and the other to test for white matter microstructural parameters. Both models also included age, sex and education as predictors. Following this, non-parametric correlations (Spearman’s Rho) were conducted between significant predictors and DDM indices, and DDM parameters and SAT to explore the directionality and significance of the relationships.

## Results

### Group differences in visual acuity

No significant differences were found in visual acuity between older and younger age groups (F(1,49) = 1.239, p=.276) (Figure 3).

**Figure 3.**
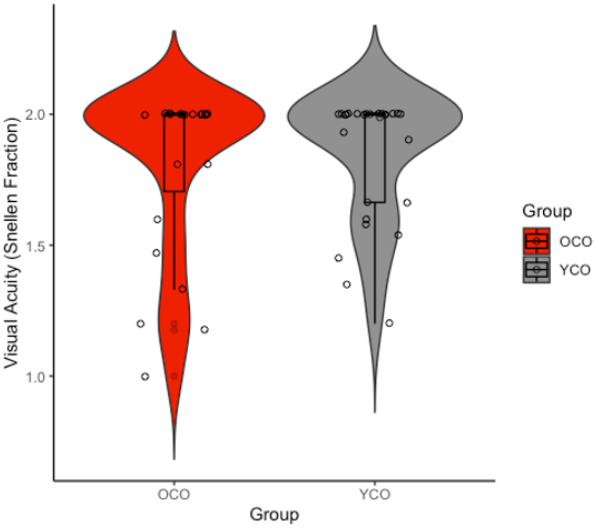
Group comparisons between older and younger adults’ visual acuity (Snellen Fraction). MANOVA results showed no significant difference in visual acuity between younger and older adults. OCO=older control participants (red), YCO=younger control participants (grey). **p<.001, *p<.05

### Group differences in RT and DDM parameters

A significant omnibus effect of group was present for DDM, RT and variance, and SAT outcome measures (F(16,24)=61.45, p=<.001; Wilk’s Λ = 0.215, partial η^2^=.785). SAT (F(1,49)=40.06, p<.001) and mean RT (F(1,49) = 12.133, p=<.001) were significantly higher in older adults than younger adults. No significant group differences were present in variance (F(1,49) = 1.36, p=.249). Significant group differences were present in non-decision time (t) (F(1,49)=4.857, p=.034) and boundary separation (a) (F(1,49)=5.341, p=.026), with older participants having greater values for each parameter. No group difference in drift rate was present (F(1,49)=.001, p=0.971; Figure 4).

**Figure 4.**
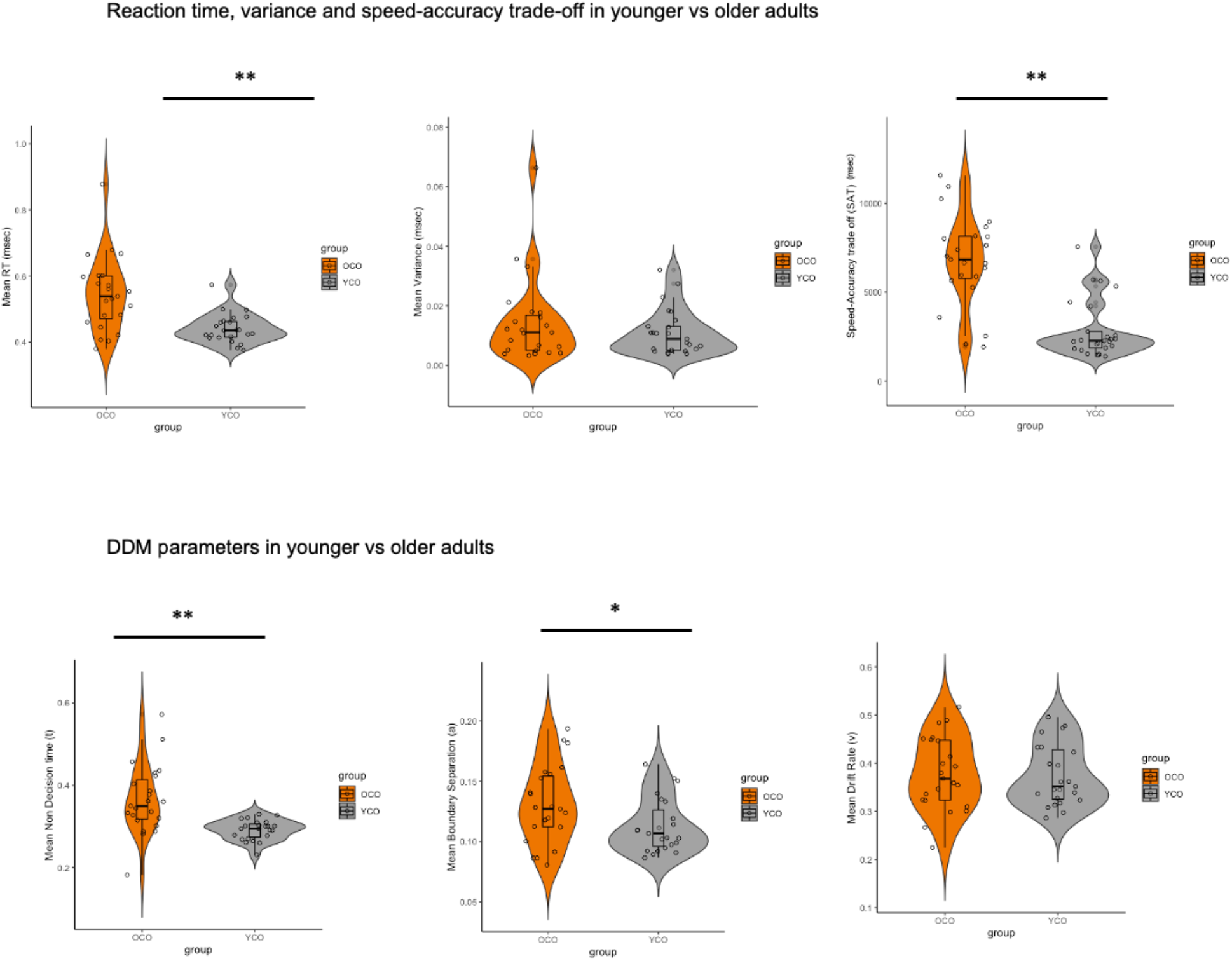
Group comparisons between older and younger adults’ mean RT, RT variance, SAT performance and DDM parameters. MANOVA results showed mean non-decision time (perceptual and motor processing) and mean boundary separation values (information threshold for decision making) are significantly greater in older adults than younger adults. OCO=older control participants (orange), YCO=younger control participants (grey). **p<.001, *p<.05

### MRI results

#### Metabolic differences between older and younger adults

Group comparisons between older and younger participants showed no significant differences in GABA levels in the ACC, OCC or the PPC (Figure 5a). Older participants had significantly lower Glx (U=189, p=.004) and NAA (U=109, p<.001) in the ACC than younger adults. A trend towards significantly lower myoinositol in older adults in comparison to younger adults in the ACC was also observed (U=234, p=.058). In the PPC, older adults showed significantly lower NAA (U=190, p=.011), and a trend towards lower Glx (U=231, p=.054) than younger adults.

**Figure 5.**
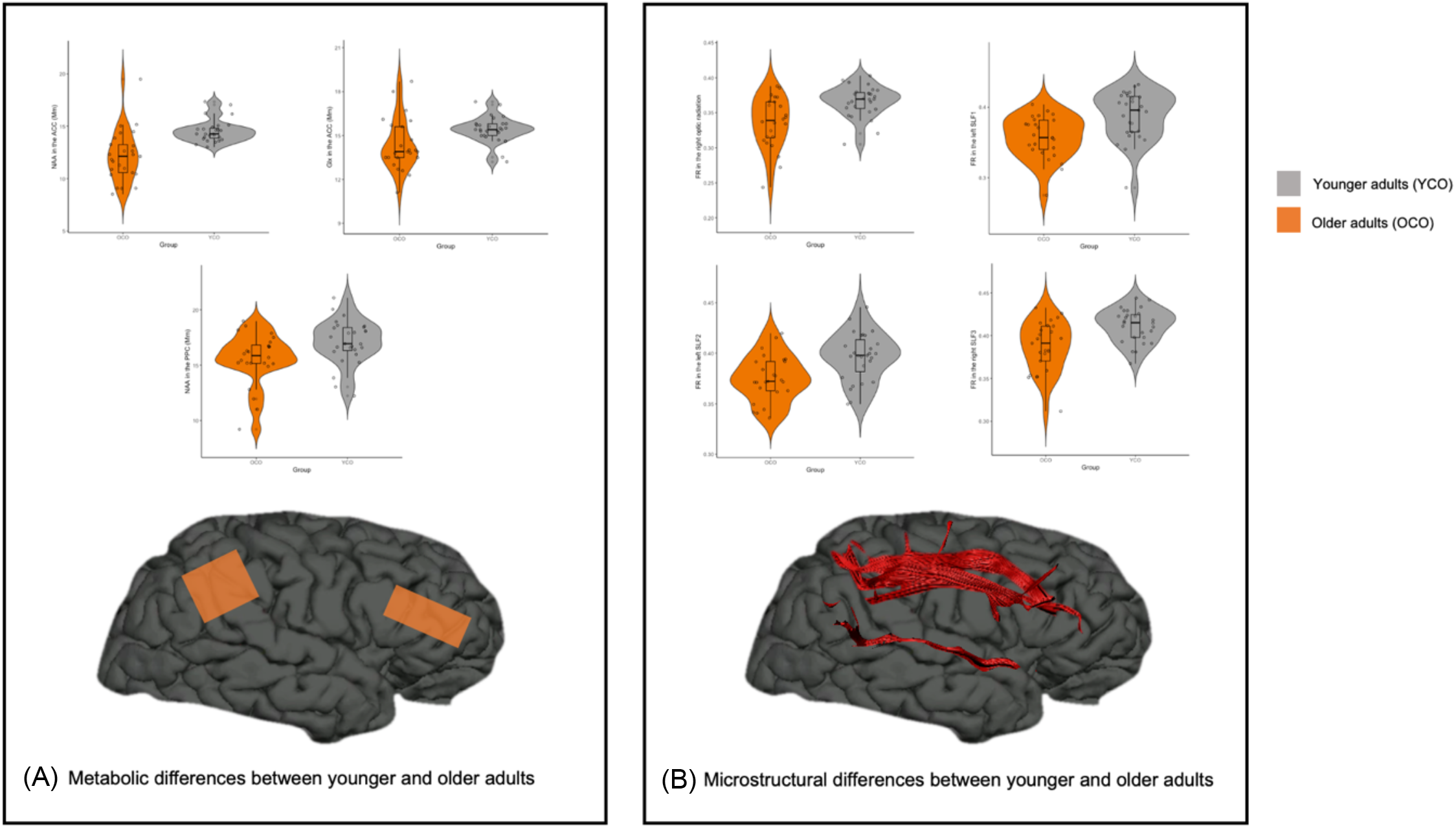
Metabolic and microstructural differences between younger and older adults. (A) Significant group comparisons for metabolites in voxels of interest between older and younger adults (B) Significant group comparisons for FR in tracts of interest between older and younger adults. FR was significantly lower in the fornix, optic radiation, SLF1, 2 and 3 in older adults in comparison to younger adults. OCO = older adults, YCO = younger adults **p<.001, *p<.0.05

#### Diffusion weighted MRI differences between younger and older adults

Significantly lower restricted fraction was shown in the older control group in the fornix (U=75, p<.001), right optic radiation (U=154, p=.001), left SLF1 (U=152, p=.001), left SLF2 (U=170, p=.002) and right SLF3 (U=169, p=.002) (Figure 5b).

Significantly higher FA in the fornix (U=22, p<.001), right optic radiation (U=207, p=.027), left ILF (U=203, p=.014), right ILF (U=157, p=.001), left SLF1 (U=131.5, p<.001), left SLF2 (U=195, p=.009), and right SLF2 (U=218, p=.029), and right SLF3 (U=163, p=.001) was found in younger adults in comparison to older adults. Significantly higher MD in the older group in comparison to the younger control group was found in the fornix (U=84, p<.001), left optic radiation (U=78, p<.001), right optic radiation (U=131, p<.001), right ILF (U=203, p=.014), right SLF1 (U=175, p=.003), right SLF3 (U=175, p=.003). Radial diffusivity was significantly higher in the older control group in all tracts of interest except the right SLF1: fornix (U=39, p<.001), left optic radiation (U=145, p=.001), right optic radiation (U=139, p<.001), left ILF (U=189, p=.007), right ILF (U=143, p<.001), left SLF1 (U=130, p<.001), left SLF2 (U=256, p=.199), right SLF2 (U=178, p=.003), left SLF3 (U=197.5, p=.010), right SLF3 (U=135.5, <.001) (Figure 5). Significantly greater axial diffusivity in older adults was found in the fornix (U=529, p<0.001), and significantly lower axial diffusivity in older adults was found in the SLF1 left (U=178, p=0.005).

### Neural piedictois of DDM parameters

Mean DDM parameters were correlated with SAT to determine how these relationships differed between younger and older groups. In younger adults, SAT was positively correlated with mean boundary (rho=.412, p=.041), and mean non decision time (rho=.388, p=.049) but not mean drift rate (rho=-.373, p=.067). In older adults, SAT was significantly correlated with boundary separation (boundary: rho=.322, p=.04) but not drift rate or non-decision time (drift: rho=.130, p=.535; non-decision: rho=.196, p=.381) (Figure 6a). Three separate linear regression models were conducted that entered age, sex and education and all metabolite measurements as predictors to assess how much variance they accounted for in the three DDM parameters of non-decision time, boundary separation and drift rate. Variation in nondecision time (t) was significantly predicted by age in the first linear regression model (adj R_2_ = 0.254, beta=0.519, p<.001) and by NAA in the ACC in the second model (adj R_2_=0.058, beta=-0.278, delta R_2_=0.09, p=0.048). Age also significantly predicted boundary separation value (a) (adj R_2_=0.410, beta=0.246, p=0.008) but no metabolites added to the fit of the model significantly. Finally, no entered predictors were shown to significantly predict drift rate. The same linear regression models as described above were also used to test for microstructural parameters as predictors together with age, sex and education. Non decision time (t) was seen to be significantly predicted by MD in the left SLF 3 (adj R_2_=0.326, beta=0.553, p=.004), MD in the SLF 2 right (adj R_2_=0.436, beta=0.484, p<.001), MD in right optic radiation (adj R_2_=0.529, beta=0.496, p=.002), and RD in left SLF 3 (adj R_2_=0.297, beta=0.394, p=.004). Boundary separation (a) was significantly predicted by age (adj R_2_=0.101, beta=0.437, p=.002), MD in left optic radiation (adj R_2_=0.070, beta=-0.763, p<.001), FR in the fornix (adj R2=0.058, beta=-0.836, p<.001), FR in right ILF (adj R_2_=0.342, beta=0.581, p=.002) and L1 in left optic radiation (adj R_2_=0.131, beta=-0.540, p<.001). Drift rate was predicted by FA in the left ILF (adj R_2_=0.062, beta=-0.468, p=.005), MD in left SLF 3 (adj R_2_=0.058, beta=.498, p=.008), L1 in the right ILF (adj R_2_=.048, beta=.548, p=.001), and L1 in left ILF (adj R_2_=.200, beta=-0.314, p=.026), but age did not contribute to the model.

## Discussion

Healthy aging leads to response slowing and increased speed-accuracy-trade-offs (SAT) but the cognitive components and neural mechanisms contributing to these changes remain poorly understood. The present study therefore aimed i) to characterise age-related differences in cognitive components contributing to response slowing using the EZ DDM model, ii) to identify age-related differences in metabolic and microstructural properties of key structures within the visual perceptual and attention networks using advanced MRI and MRS, and iii) to explore how these brain measurements were linked to age-related differences in SAT and DDM parameters of decision-making processes in a modified ANT flanker task.

Consistent with the literature we found that older adults without reduced visual acuity showed longer SAT as well as greater non-decision time and boundary separation values compared with younger adults. This pattern of results suggests a slowing of perceptual and/or motor elements of RT as reflected in larger non-decision time, as well as a higher boundary of information to be accrued before a decision can be reached, as reflected in higher boundary separation values. An emphasis on accuracy in place of faster speed has previously been shown to result in increased boundary separation in visual perceptual tasks in younger (Zhang & Rowe, 2014) and older adults (Starns & Ratcliff, 2010), suggesting a close link between age-related differences in the pattern of DDM parameters and the typical age-related shift in SAT.

We also demonstrated that older adults show reduced levels of Glx and NAA in the ACC and lower levels of NAA in the PPC compared with younger adults. NAA and Glx are both markers of neuronal metabolism (Newsholme et al., 2003) and are considered to play a key role in energy metabolism in neural mitochondria (Lu et al., 2004). These findings are consistent with accumulating evidence of energy depletion as a key component of biological aging (Raz & Daugherty 2018). With normal aging, the accumulation of biological ‘imperfections’ such as protein aggregation are thought to impair mitochondrial function and cause low-level inflammation, resulting in reduced glucose uptake, synaptic deterioration, and gliosis (Currais, 2015), and in turn to further energy reduction. The here observed reductions in NAA and Glx in the ACC in older adults may reflect a decreased neuronal metabolism due to the accumulation of these biological ‘imperfections’. This suggests that aging affects neuronal metabolism in frontal and parietal attention regions but not in the OCC, consistent with some evidence suggesting that healthy aging is particularly associated with a reduction in mitochondrial energy metabolism in frontal brain regions (Reddy & Beal, 2008, Reutzel et al., 2020, Yin et al., 2014).

Our tractography analyses provided evidence for lower white matter FR and FA and greater MD, RD and L1 in older compared to younger adults. Microstructural differences were observed in all white matter pathways of interest, i.e., the optic radiation, the ILF, the SLF and the fornix. Overall, this pattern of widespread age-related differences in white matter microstructure is consistent with the literature and suggests that all of the studied white matter connections of visual perception and attention networks are detrimentally affected by age. While previous studies into age-related differences in white matter microstructure have primarily focused on DTI measurements and have generally reported reduced FA and increased MD and RD in pathways including the ILF (Inano et al., 2011), SLF (Lai & Wu, 2014) and the fornix (Metzler-Baddeley et al., 2011), fewer studies have applied the restricted diffusion signal fraction FR from CHARMED. Reductions in FR suggest a decrease in the density of axons, that may occur due to age-related loss of myelin and/or axonal loss secondary to Wallerian degeneration. Although DTI and FR measures were corrected for partial volume effects with the Free Water Elimination Method (Pasternak et al., 2009), it cannot be completely ruled out that atrophy-induced free water contamination may have biased these indices particularly in regions susceptible to partial volume contamination such as the fornix (Metzler-Baddeley et al 2012; see Parker et al., 2021).

Further we sought to determine which brain measurements predicted DDM parameter performance in younger and older adults. Consistent with our hypotheses, non-decision time was predicted by differences in SLF and optic radiation microstructure in older adults and NAA in ACC in younger adults. Thereby larger diffusivity in the SLF was associated with longer non-decision times and this was particularly driven by older adults.

Non-decision time has previously been linked to fronto-parietal fMRI activity in older adults, therefore these observed findings complement and extend these results (Madden et al., 2020). Similarly, age-related increases in variability and reductions in visual functions requiring the suppression of irrelevant perceptual information were associated with individual variability in fronto-parietal white matter microstructure (Chadick et al., 2014). Thus, age-related deterioration in the SLF - a white matter pathway crucial for the effective communication within the fronto-parietal attention network and hence between sensory and motor networks - is likely to contribute to the overall slowing of sensory and/or motor processing components reflected in the non-decision time. In addition, it has also been documented that older adults show reduced functional connectivity in fronto-parietal networks and that this is related to increased distractibility and diminished attentional focus with age (Campbell et al., 2012, Madden et al., 2020). Thus, the observed pattern of associations between age-related variation in non-decision time and SLF microstructure may reflect age-related difficulties in perceptual suppression or inhibition (Chadick et al., 2014) and/or a greater reliance in top-down processing as a response to diminished low-level sensory input (Lai et al., 2020). As top-down processing is thought to occur in the dorsal processing stream in frontal and parietal brain regions (Gilbert & Sigman, 2007), non-decision time may capture elements of both bottom-up and top-down perceptual changes in aging that are mediated by the dorsal attention processing networks. These results suggest that older adults attempt to rely on top-down processing - due to fronto-parietal tracts showing significant relationships with non-decision time - but that these connections have lower WM integrity and thus result in longer non-decision times and overall lengthened SAT. This interpretation of results is in line with previous findings of age-related mediation of the relationship between frontoparietal activation and impaired top-down attention (Madden et al., 2014).

Further we observed a contribution of NAA in the ACC to the variation in non-decision time. As NAA is thought to be a marker for neuronal health and integrity, and the ACC is linked to inhibitory functioning as it is thought to be part of the anterior salience network responsible for conflict monitoring (Dosenbach et al., 2006). Thus it is possible that age-related reductions in neuronal health in the ACC may contribute to longer non-decision time on the basis of poorer inhibition. A relationship between NAA in the ACC and cognitive inhibitory processes has previously been observed (Grachev et al., 2001), suggesting that NAA in the ACC may play a role in perceptual interference which may be represented by non-decision time. However, as non-decision time represents both perceptual and motor aspects of the RT, and NAA in the ACC has also been linked to motor inhibition processes (Leland et al., 2008; Paus, 2001; Rubia et al., 2001), it remains unclear which element of non-decision time may be predicted by metabolite changes in the ACC.

Our results have also shown that beyond age, boundary separation was significantly predicted by microstructural differences in the optic radiation, ILF and fornix. These findings are consistent with the view that noisy sensory processing due to age-related impairments in the microstructural integrity of white matter pathways important for bottom-up perceptual processing will influence the criterion threshold for decision making. In other words, older participants faced with noisier sensory input, as reflected by higher non-decision time values, will become more cautious and will increase their boundary threshold before eliciting a response, consistent with the age-related shift in SAT.

Drift rate did not significantly differ between younger and older adults; however, our results showed it to be predicted by FA and L1 in the ILF and MD in the SLF3, suggesting that the rate of evidence accumulation through the perceptual system is critically dependent on the microstructural integrity of white matter connections that propagate that information. Previous evidence suggests that drift rate was related to attentional allocation (Madden et al., 2010, 2020) and the SLF 3 is thought to be pivotal for the allocation of visuo-spatial attention (De Schotten et al., 2011). Moreover, information processing speed, to which drift rate is closely related, is associated with temporal lobe oscillations (Chauvière, 2020) and stimulus-related attention has been associated with neural activity in the inferior temporal cortex (Zhang et al., 2011). This may explain why microstructure in the ILF was shown to predict drift rate in this study.

### Conclusions

In summary, this study replicated findings of age-related increases in SAT, non-decision time and boundary separation and provided novel evidence of age-related reductions in Glx and NAA in the ACC and PPC, suggestive of impaired neuronal health in these regions. Older adults also showed widespread white matter microstructural impairments manifest as lower FA and FR and greater MD, RD and L1 in all tracts of interest that may reflect age-related loss of myelin, axonal packing and atrophy. Importantly we demonstrated that performance in non-decision time elements of RT processing was predicted by microstructure measurements in the SLF and optic radiation as well as by NAA in ACC, while boundary separation was predicted by differences in optic radiation, ILF and fornix. These findings suggest that both bottom-up and top-down processes contribute to response slowing in aging. More specifically, noisy sensory and perceptual input due to impairments in the microstructural integrity of white matter in pathways mediating bottom-up processing that may not be compensated by top-down predictions due to a decline in neuronal health in anterior attention regions (ACC), may lead to an increase in boundary separation or criterion threshold of decision-making in older adults and hence an increase in SAT.

